# A novel role for Neurog2 in MYCN driven neuroendocrine plasticity of prostate cancer

**DOI:** 10.1101/2024.08.16.607954

**Authors:** Prachi Walke, Jared D.W. Price, Frederick S. Vizeacoumar, Nickson Joseph, Vincent Maranda, Bari Chowdhury, Jay Patel, Yue Zhang, He Dong, Lara New, Ashtalakshmi Ganapathysamy, Li Hui Gong, Hussain Elhasasna, Kalpana K. Bhanumathy, Yuliang Wu, Andrew Freywald, Anand Krishnan, Franco J. Vizeacoumar

## Abstract

Neuroendocrine prostate cancer (NEPC) presents a formidable clinical challenge owing to its aggressive progression and resistance to conventional therapies. A key driver of NEPC is the overexpression of *MYCN*, a well-established oncogene associated with neuroendocrine tumors. However, efforts to directly inhibit the N-Myc protein encoded by this gene have resulted in limited success, thereby hindering therapeutic advancements. To overcome this obstacle, we conducted unbiased genome-wide screening using isogenic prostate cancer cell lines to identify the synthetic vulnerabilities of *MYCN*. Among the identified candidates, *NEUROG2* emerged as a significant candidate. Neurog2 is a proneural transcription factor (PTF) known for its role in developmental processes and trans-differentiation of adult cells. Our findings demonstrate that Neurog2 depletion does not affect non-malignant cells, but significantly suppresses the growth of *MYCN*-overexpressing cells and tumors in orthotopic NEPC models. Furthermore, our observations indicate that the Neurog2-mediated regulation of PTFs can facilitate NEPC development. Thus, targeting Neurog2 holds promise as an effective therapeutic strategy for *MYCN*-overexpressing NEPC.

## Introduction

Prostate cancer (PC) is a common disease in the male population, exerting a profound impact on the public health systems worldwide^1^. Despite advances in diagnostic techniques and therapeutic modalities, managing advanced PC subtypes continues to present clinical hurdles. Androgen deprivation therapy (ADT), which reduces testosterone levels to castration levels, is the cornerstone treatment for primary prostate adenocarcinoma in patients with inoperable tumors^2–4^. However, the emergence of castration-resistant prostate cancer (CRPC) after ADT underscores the urgent need for additional therapeutic strategies^5^. The transition from adenocarcinoma to CRPC is orchestrated by several mechanisms, including intratumoral androgen synthesis, mutation and overexpression of androgen receptor (AR), and upregulation of AR coactivators, perpetuating AR signaling despite castration levels of testosterone^5,6^. While new-generation anti-androgen therapies have demonstrated some improvements in the overall survival of CRPC patients, therapeutic resistance and development of neuroendocrine prostate cancer (NEPC) remain major challenges, necessitating continued exploration of novel therapeutic avenues^4,7–10^.

Among the spectrum of PC subtypes, NEPC has emerged as particularly aggressive, characterized by treatment resistance and rapid disease progression. Although NEPC may develop *de novo*, it is frequently triggered by hormonal (anti-androgen) therapies, resulting in dismal prognosis with limited therapeutic options^4,7^. Molecular alterations, including amplification and overexpression of the *MYCN* oncogene, frequently accompany the transition from PC to NEPC, underlining the pivotal role of *MYCN* and the N-Myc transcription factor it encodes in driving this aggressive PC subtype^9,11,12^.

The pursuit of therapeutic interventions targeting N-Myc has witnessed significant strides in recent years, despite its initial classification as an ’undruggable’ molecule. Various strategies, including small molecules targeting the N-Myc-MAX interaction or direct modulation of N-Myc function, have shown promise in preclinical models^13^. Additionally, innovative approaches such as therapeutic mini-proteins and targeted degradation strategies offer novel avenues for inhibiting N-Myc activity. Encouragingly, the development of MRT-2359, a phase I/II clinical trial candidate that indirectly causes N-Myc downregulation, underscores the growing momentum of exploring anti-N-Myc therapies^14^. Unfortunately, despite these efforts, no therapy targeting N-Myc is currently available for clinical application.

Here, instead of targeting N-Myc, we aimed to elucidate the genetic dependencies of *MYCN* in prostate cancer cells. Specifically, we applied a genetic principle known as Synthetic Dosage Lethality (SDL), where the loss of function of a gene leads to lethality only when another gene, such as *MYCN*, is overexpressed^15,16^. This paradigm holds immense potential not only in indirectly targeting overexpressed oncogenes but also in facilitating the development of personalized medicine based on the functional status of an oncogene in individual patients. Expanding on this genetic principle, we conducted extensive genome-wide loss-of-function screens employing both sgRNA/CRISPR/Cas9 and shRNA platforms and isogenic cell line models of NEPC. Our findings revealed, for the first time, a novel genetic interaction between *NEUROG2* and *MYCN*. Subsequent *in vitro* studies have confirmed that Neurog-2 is not expressed and is dispensable for the survival of non-malignant cells. However, *in vitro* viability studies and *in vivo* studies in orthotopic tumor models suggest that the loss of function of Neurog-2 suppresses the growth of *MYCN*-overexpressing cells and tumors, respectively. Overall, our findings have promising implications for the development of targeted therapies customized for N-Myc-driven NEPC.

## Results

### Integration of CRISPR and shRNA screens identifies genetic dependencies of MYCN overexpressing NEPC cells

We explored SDL interactions of *MYCN* to identify novel targets for indirectly inhibiting N-Myc-dependent NEPC tumors. To this end, we performed loss-of-function genetic screens in an isogenic model generated by overexpressing *MYCN*. We used 22Rv1 and 22-MYC cells (N-Myc overexpressing 22Rv1, a gift from the Rickman laboratory), in which N-Myc overexpression induces poorly differentiated invasive PC that is molecularly similar to NEPC^8^ to recover *MYCN* specific SDL hits. In our experience, screens in isogenic models tend to produce fewer false positives and capture hits specific to the query gene^17–21^. Prior to screening, we confirmed the overexpression of N-Myc in 22-MYC cells by western blotting and qRT-PCR **(Figure 1A, B)**. We also confirmed the potential of 22-MYC cells to form NEPC type tumors *in vivo*, as evidenced by the elevated expression of the NED marker neuron-specific enolase (NSE) in 22-MYC-driven tumors compared to that in 22Rv1 cells **(Figure 1C)**.

**Figure 1:**
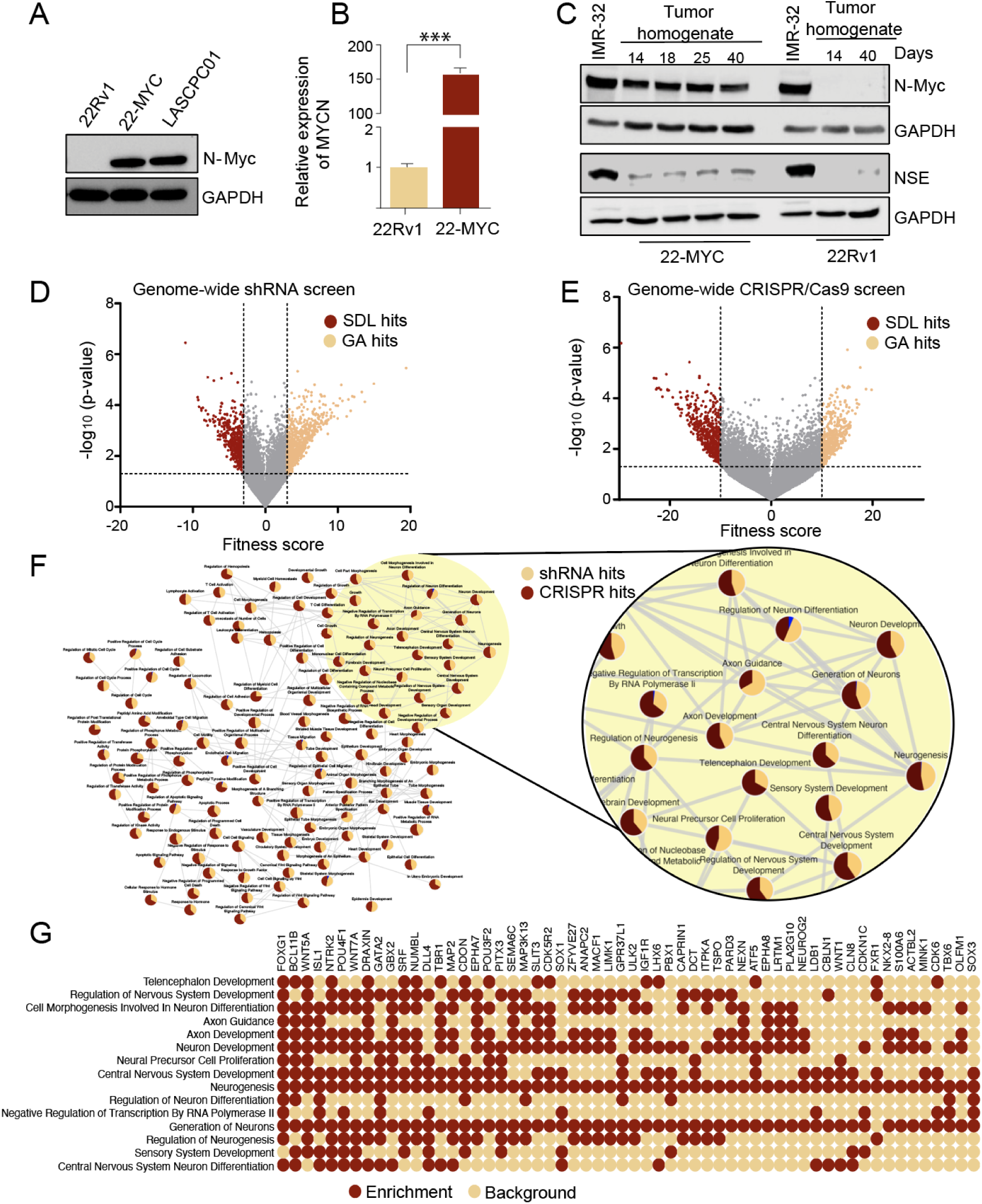
Pooled shRNA and CRISPR screens for the exploration of NEPC targets: (A) N-Myc protein levels in 22Rv1, 22-MYC and LASCPC01 cells. (B) N-Myc encoding mRNA expression in 22Rv1 and 22-MYC cells. Data presented as mean ± SE, n=3; Standard ‘t’ test; ***p<0.01. C) Protein levels of N-Myc and the NSE NED marker in 22-MYC and 22Rv1 subcutaneous tumor xenografts harvested at the indicated days. Innately N-Myc^high^ IMR-32 neuroblastoma cells were used as a positive control for N-Myc and NSE. (D-E) Volcano plots representing genome-wide screening in 22-Rv1 and 22-MYC cell line pair using pooled shRNA (left) and CRISPR/Cas9 screens (right). The red dots represent SDL hits and the orange dots on the right represent possible growth advantage (GA) hits. Grey dots represent genes whose loss of function did not significantly affect cell viability in the screens. The y-axes show the significance of the hits, and the x-axes represent the fitness score computed for each gene by comparing 22-Rv1 cells with 22-MYC cells. Cut-offs were determined based on data distribution within each screen. (F) Pathway enrichment analyses of both shRNA and CRISPR screens. GSEA was performed and genes with p≤0.005 and FDR≤0.1 were used to select enriched pathways. Each node is divided into two colors representing the percentage of genes identified in each screen. Enlarged area represents neuron-associated terms. (G) Dot plot showing the top 58 genes associated with neuronal plasticity (Dark color represents pathway enrichment, light color represents no-enrichment background).

For genome-wide SDL screening, we utilized two independent pooled screening platforms: an shRNA gene knockdown lentiviral library and a CRISPR/Cas9 gene knockout lentiviral library. These platforms targeted approximately 18,000 genes to identify SDL interactions specific to *MYCN*. The Pearson correlation coefficient showed a strong correlation (r > 0.89) between the replicas considered for each screen **(Supplementary Figure 1)**. Overall, the shRNA screen identified 581 SDL hits, whereas the CRISPR/Cas9 screen identified 553 hits, whose selective elimination selectively suppressed N-Myc-positive cells **(Figure 1D, E; Supplementary Table S2)**. Some of the promising hits identified included heat-shock proteins and related co-chaperones, such as HSPA5, HSP90B1, CCT5, HSPB6, and UBXN1, which have critical roles in prostate cancer, including combating ER stress^22,23^. Interestingly, several of the identified SDL hits, such as *CDK6, CERK, TBX2,* and *MBD3* were also identified in another genome-wide screen performed for *MYCN*-driven neuroblastoma, which is another cancer of neuroendocrine origin^24^, suggesting that some of our hits might also be applicable to other *MYCN*-driven malignancies. Alternatively, this might also mean that some of our SDL hits may be relevant to the neuroendocrine tumor (NET) phenotype *per se.* Specifically, among the SDL hits, the shRNA screen identified proneural transcription factors (PTFs), such as *NEUROG2* and *POU3F2,* along with transcriptional targets of N-Myc, including *PKMYT1, CKS1B, RBP5,* and *CHEK2*^25–27^. Similarly, CRISPR/Cas9 screening identified several neuromodulators, including *SOX17, FOXG1, PITX3, ONECUT2,* and *GBX2*, which favor NED and potentially contribute to NEPC. Although the overlap between the two screens was anticipated to be modest^28,29^ amalgamating data from both platforms revealed several genes operating within the shared pathways **(Figure 1F)**. Notably, we identified genes associated with axon guidance (FDR<0.0001), Wnt signaling (FDR<0.0001), neurogenesis (FDR<0.0001), neuron development (FDR<0.0001), and sensory system development (FDR<0.0001), all of which are significantly relevant to neuroendocrine tumors, including NEPC^30–34^. Particularly noteworthy from the enrichment analyses were approximately 58 genes implicated in neuronal plasticity, a trait closely associated with NEPC **(Figure 1G)**. Thus, the independent identification of NED- and NEPC-relevant hits in both shRNA and CRISPR/Cas9 screens underscores the robustness and advantages of the dual-screening strategy employed in this study.

### Coupling differential essentiality with differential expression to prioritize SDL hits

To evaluate the significance of SDL genes specific to NEPC, we explored the genes differentially expressed between CRPC and NEPC patient samples using publicly available data^35^. Among the screened hits, 323 genes exhibited significantly increased expression in NEPC samples compared to CRPC samples. Unsupervised clustering of these 323 genes revealed distinct gene expression patterns between NEPC and CRPC **(Figure 2A)**. Notably, several genes related to neuronal plasticity, including *KIT, FOXG1, GBX2, WNT4, SOX1, ONECUT2, NEUROG2* and *ISL1*, were clustered together. This indicates that the combined evaluation of differentially expressed genes and SDL candidates is a powerful approach to identify potential targets that regulate NEPC biology^36–43^.

**Figure 2:**
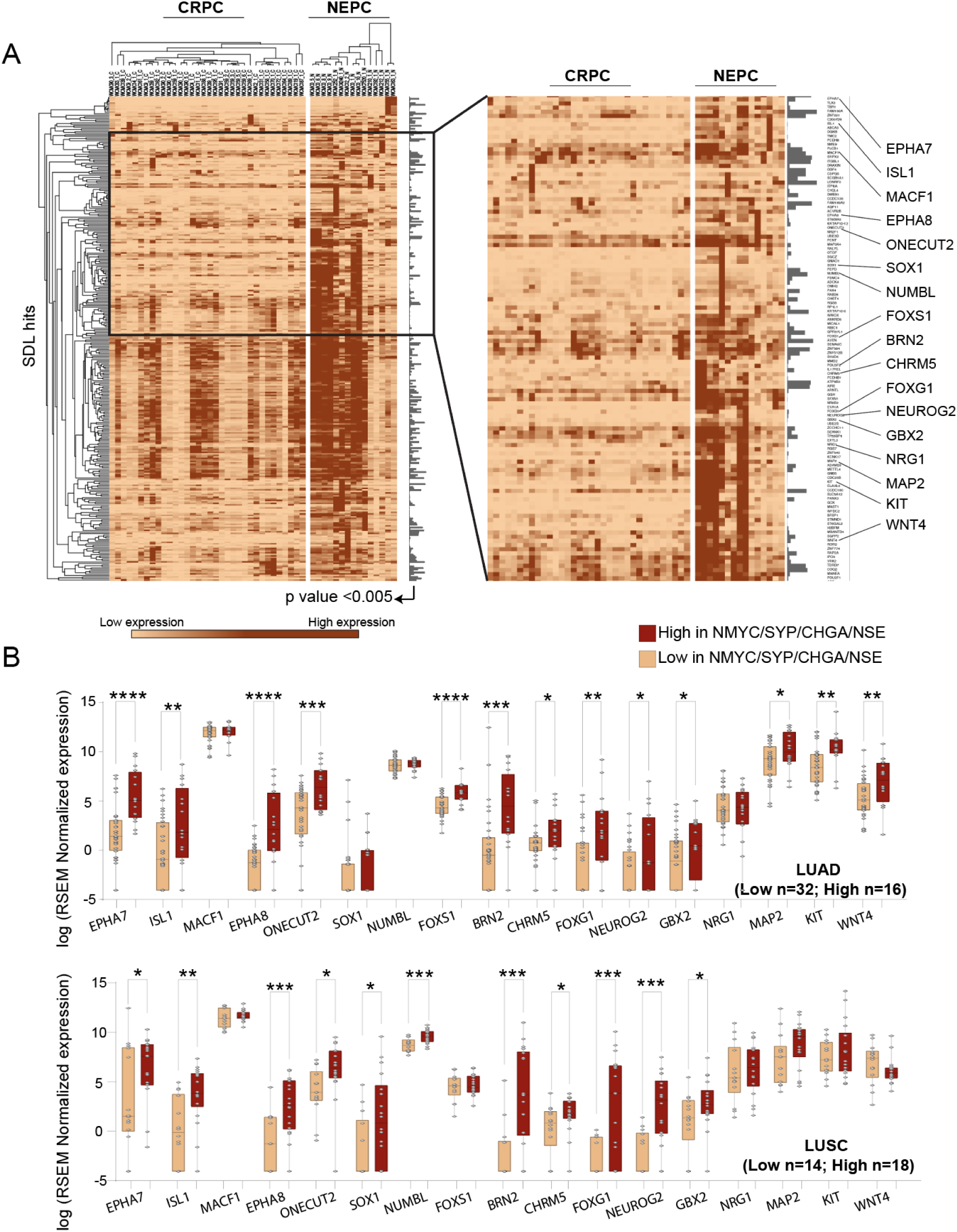
Prioritization of hits using patient data: (A) Unsupervised Pearson clustering of gene expression profiles for prostate cancer patient samples from Beltran et al. (Nat. Methods, 2016), highlighting 323 genes with a significant difference (Wilcoxon rank-sum test, p<0.005) between CRPC and NEPC. The inset image magnifies genes exhibiting the most pronounced differential expression. (B) Expression analysis of lung cancer patient samples from TCGA, illustrating significant differences (one-tailed unpaired t-test: *p<0.05; **p<0.01; ***p<0.001; ****p<0.0001) in the 17 genes associated with neuronal plasticity, after classification based on the expression of the *NMYC/SYP/CHGA/NSE* signature. Top panel represents lung adenocarcinoma (LUAD) and bottom lung squamous cell carcinoma (LUSC).

To further validate these findings, we examined The Cancer Genome Atlas (TCGA) to assess the expression patterns of some of these genes in lung squamous cell carcinoma (LUSC) and lung adenocarcinoma (LUAD), as they can potentially transition to small cell lung cancer that shares more neuroendocrine features^44^. The cellular plasticity of both tumor types has the potential to transform them into neuroendocrine tumours^45,46^. Therefore, to compare the expression status of SDL hits within these tumor types, we categorized patient data based on the expression of *MYCN* and NED markers such as SYP, CHGA, and NSE. As noted above, our analysis revealed that several SDL candidates, including *ISL1, NEUROG2, EPHA7, EPHA8, ONECUT2, POU3F2, CHRM5, FOXG1 and GBX2* were significantly overexpressed in LUAD and LUSC cases with neuroendocrine features, represented by high expression of *MYCN* and NED markers **(Figure 2B)**. These findings not only highlight the potential role of neuronal plasticity genes in NEPC and *MYCN*-overexpressing lung cancer but also highlight their broader significance in neuroendocrine tumors.

### Neurog2 is essential for the survival of N-Myc overexpressing cells

We next chose to validate two SDL hits, *NEUROG2* and *ISL1* from among the major hits, as they are both involved in neurogenesis^47,48^ and our analyses indicated that they are significantly upregulated in NEPC **(Figure 2A)**, while their therapeutic potentials and links to neuroendocrine tumors were not investigated. To determine their therapeutic potential, we initially assessed the effects of their knockout in the colony formation assay. This revealed a significantly stronger reduction in colony formation in 22-MYC cells than in 22Rv1 cells following individual knockouts of *NEUROG2* and *ISL1* **(Figure 3A)**. The differential selectivity in N-Myc-overexpressing cells validated the identification of these candidates in our genome-wide screening. To eliminate cell line-specific effects in our observations, we also examined the anti-proliferative effect of the target knockdown in N-Myc^+^/myrAKT1^+^ LASCPC-01 cells that demonstrated NEPC features using a resazurin assay and found that individual knockdown of *NEUROG2 and ISL1* effectively suppressed cell viability **(Figure 3B)**. We also confirmed the efficiency of target knockdown in LASPC-01 cells using qRT-PCR **(Figure 3C)**^49^.

**Figure 3:**
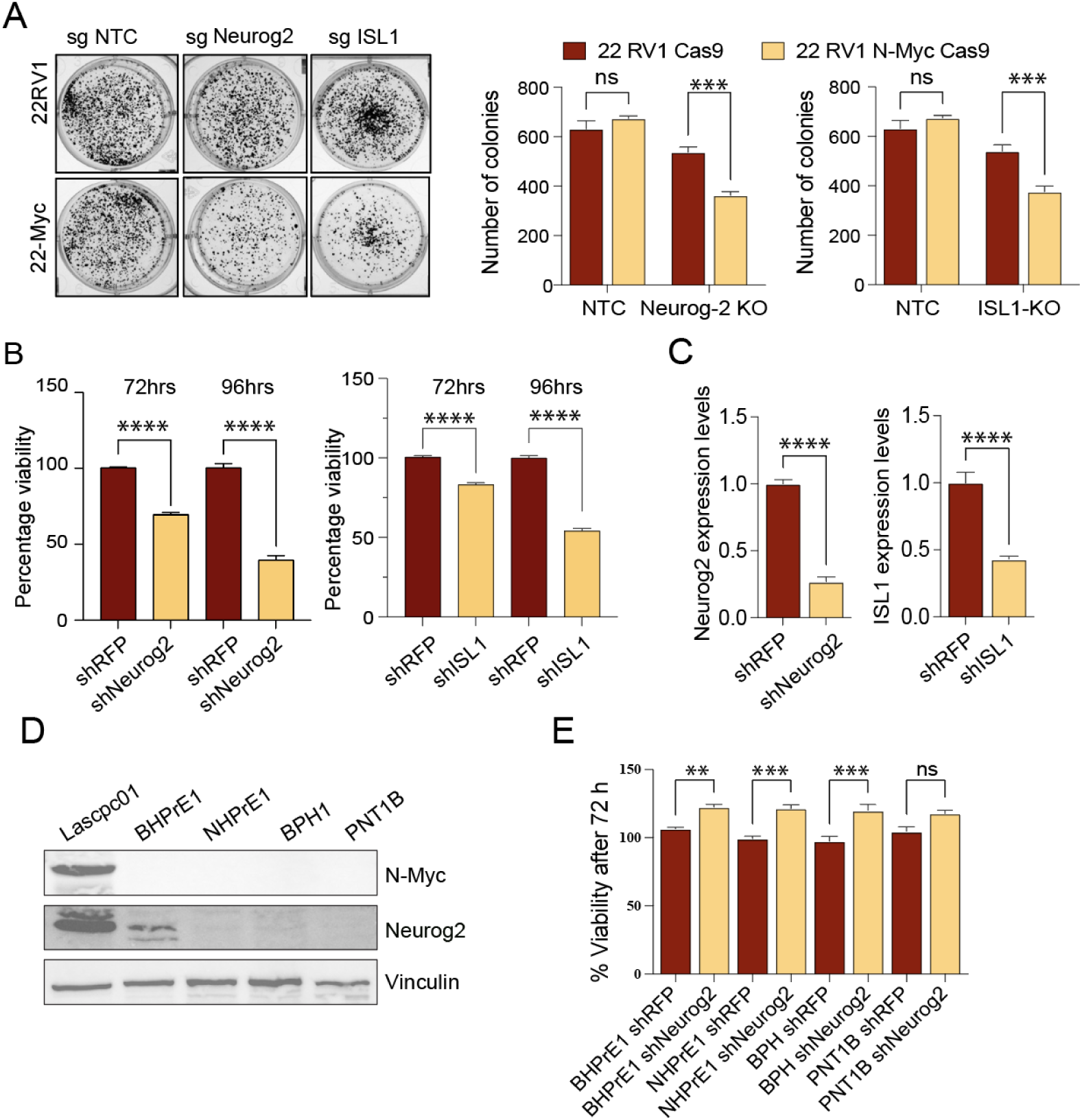
Validation of selected SDL candidates: (A) Colony formation assay and quantitation using 22-Rv1 and 22-MYC cells after NEUROG2 and ISL1 knockouts Data presented as mean ± SE n= 4; Two-way ANOVA with Sidak’s multiple comparisons test; ***p<0.001. The same NTC data was used to compare with NEUROG2 and ISL1. (B) Evaluation of cell viability using resazurin assay in LASCPC-01 cells after NEUROG2 and ISL1 knockdown. Percentages of relative viability in knockdown compared to control shRNA transduced cells are shown at the indicated time points Data presented as mean± SE; n=4; Standard ‘t’ test; ****p<0.0001. The same shRFP was used to compare with shNeurog2 and shISL1. (C) mRNA levels show efficient knockdown of NEUROG2 and ISL1 in LASCPC-01 cells, compared to non-targeting shRFP controls. Data presented as mean± SE; n=9; Standard ‘t’ test; ****p<0.0001. (D) Western blot showing protein levels of N-Myc and Neurog2 in non-malignant prostate cells. Vinculin serves as a loading control. (E) Evaluation of cell viability using resazurin assay in non-malignant cells (BHPrE1, NHPrE1, BPH1 and PNT1B) following NEUROG2 knockdown. Data presented as mean± SE; n≥ 3; One-way ANOVA with Tukey’s multiple comparisons test; **p<0.01, ***p<0.001.

Our aim was to develop targeted therapies for NEPC that do not impair normal tissue function. In line with this, it is interesting to note that we found *NEUROG2* but not *ISL1* not to be expressed in most of the normal tissues, including prostate, as assessed using the tissue and cell-specific gene expression dataset from Genotype-Tissue Expression (GTEx) Portal **(Supplementary Figure 2)**. We also examined the innate expression of Neurog2 in non-malignant prostate cell lines BPH1, BHPrE1, NHPrE1, and PNT1B and, consistent with our GTEx analyses, found that its abundance is very minimal to undetectable in these cells **(Figure 3D)**. As expected, *MYCN* was not expressed in non-malignant cells **(Figure 3D)**. Furthermore, we examined whether *NEUROG2* knockdown impaired the viability of these non-malignant prostate cells, including BPH1 cells that displayed at least minimally detectable Neurog2 expression, and found that it did not impair their viability (**Figure 3E)**. Altogether, these observations indicate that Neurog2 is dispensable to non-malignant cells but is selectively essential for the survival of *MYCN* overexpressing cancer cells.

### Neurog2 plays a key role in neuroendocrine plasticity of prostate cancer cells

NED is a hallmark of NEPC. Therefore, we examined the expression of NED markers in LASCPC-01 cells after *NEUROG2* and *ISL1* knockdown to evaluate their effect on suppressing NEPC features^50–52^. Unlike *ISL1* knockdown, *NEUROG2* knockdown suppressed NED characteristics, as evidenced by the downregulation of all three NED markers tested: SYP, CHGA, and NSE **(Figure 4A).** Similar results were observed at the mRNA level **(Figure 4B)**. These results were further corroborated by identifying positive correlations in gene expression between NED markers and Neurog2 in NEPC but not in CRPC **(Figure 4C)**. Specifically, while the correlation in gene expression between NEUROG2 and the NED markers appeared poor in CRPC (r=0.004, p=0.9 for Neurog2 vs SYP; r=0.06, p=0.71 for Neurog2 vs CHGA; r=0.006, p=0.21 for Neurog2 vs NSE), the correlation significantly improved in NEPC (r=0.72, p= 0.003 for Neurog2 vs SYP; r=0.33, p=0.21 for Neurog2 vs CHGA; r=0.45, p=0.09 for Neurog2 vs NSE), with the exception of CHGA, although its correlation also improved **(Figure 4C)**. These results indicate that, unlike *ISL1*, *NEUROG2* plays a critical role in NEPC transformation.

**Figure 4:**
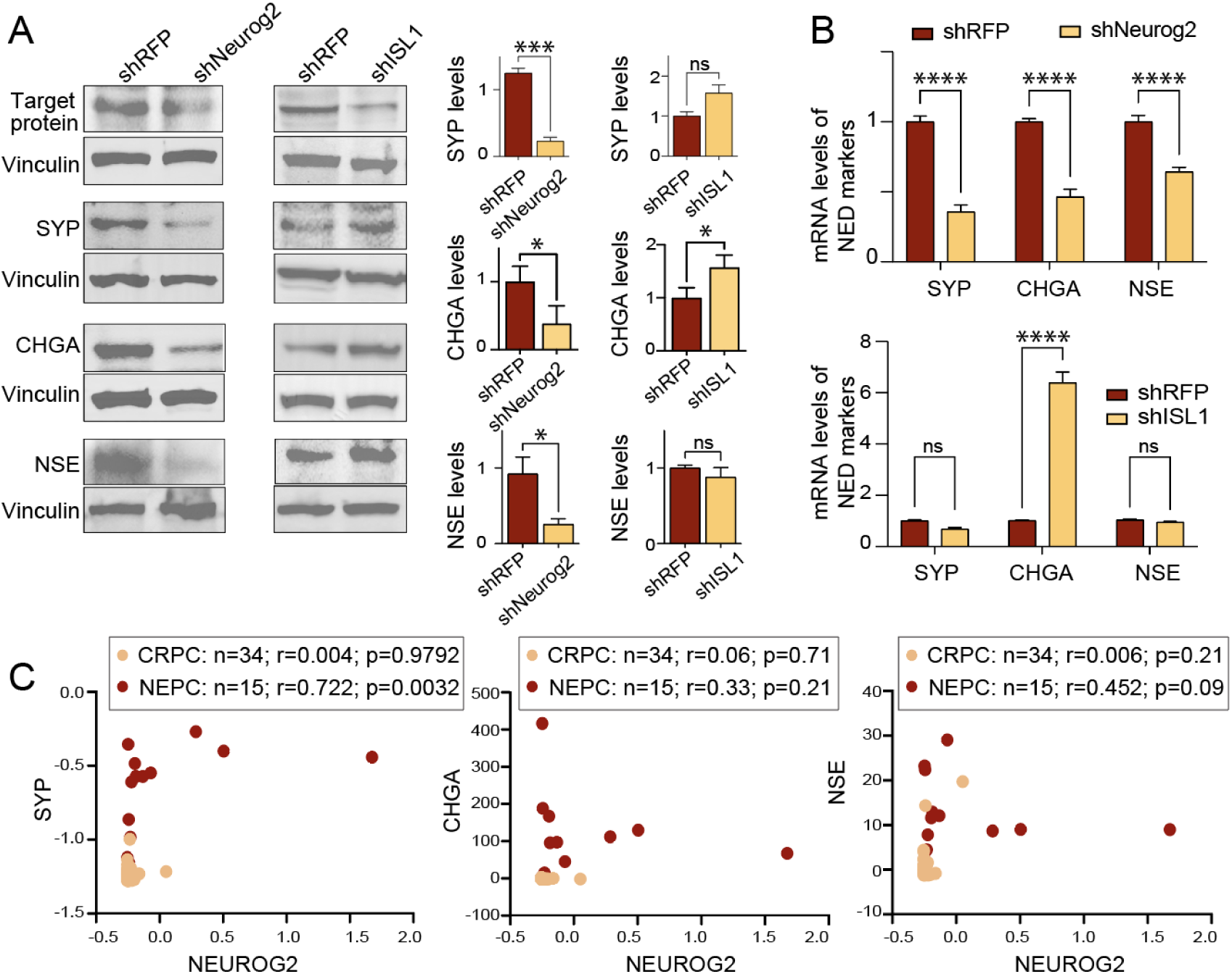
Identification of Neurog2 as a priority candidate: (A) Reduction of protein levels of NED markers (SYP, CHGA, and NSE) in LASCPC-01 NEPC cells in response to Neurog2 knockdown compared to ISL1 knockdown. Western blot images and corresponding quantifications are shown. Data represented as mean ± SE; n=3 except for NSE where n=2; Standard ‘t’ test; *p<0.05, **p<0.01, ***p<0.001 (B) Expression of NED markers (SYP, CHGA, and NSE) in LASCPC-01 cells after Neurog2 knockdown compared to ISL1 knockdown. Data presented as mean ± SE; n=9 except for NSE where n=8; Two-way ANOVA with Tukey’s multiple comparisons test; ****p<0.0001. For SYP, the same shRFP data was used to compare in both graphs. (C) Expression correlation between Neurog2 and NED markers (SYP, CHGA and NSE) in CRPC and NEPC samples from Beltran et al. (Nat. Methods, 2016).

Previous studies indicated that PTFs promote NEPC. For example, Brn2 (also known as Pou3f2) facilitates the transition from PC to NEPC^53^. Similarly, Ascl1, another PTF, has been linked to NEPC development^54^. In addition, INSM1 and NeuroD1 have also been found to contribute to NEPC^55–58^. Small cell lung cancer (SCLC), which shows features of neuroendocrine cancers, is characterized by a molecular signature that closely resembles NEPC^45^. Of note, Ascl1, NeuroD1, Pou2f3, and the transcriptional regulator Yap1 have been suggested to be critical players in SCLC, with Ascl1 and NeuroD1 recognized as potential promoters and Pou2f3 and Yap1 recognized as potential suppressors^59^. Strikingly, we found that *NEUROG2* knockdown in LASCPC-01 cells downregulated *BRN2*, *ASCL1*, *INSM1*, and *NEUROD1*, while upregulating *TAZ1,* a transcriptional co-activator with functions similar to Yap1^60,61^,indicating that functions of Neurog2 in *MYCN* expressing cells may be directed towards promoting neuroendocrine plasticity **(Figure 5A)**.

**Figure 5:**
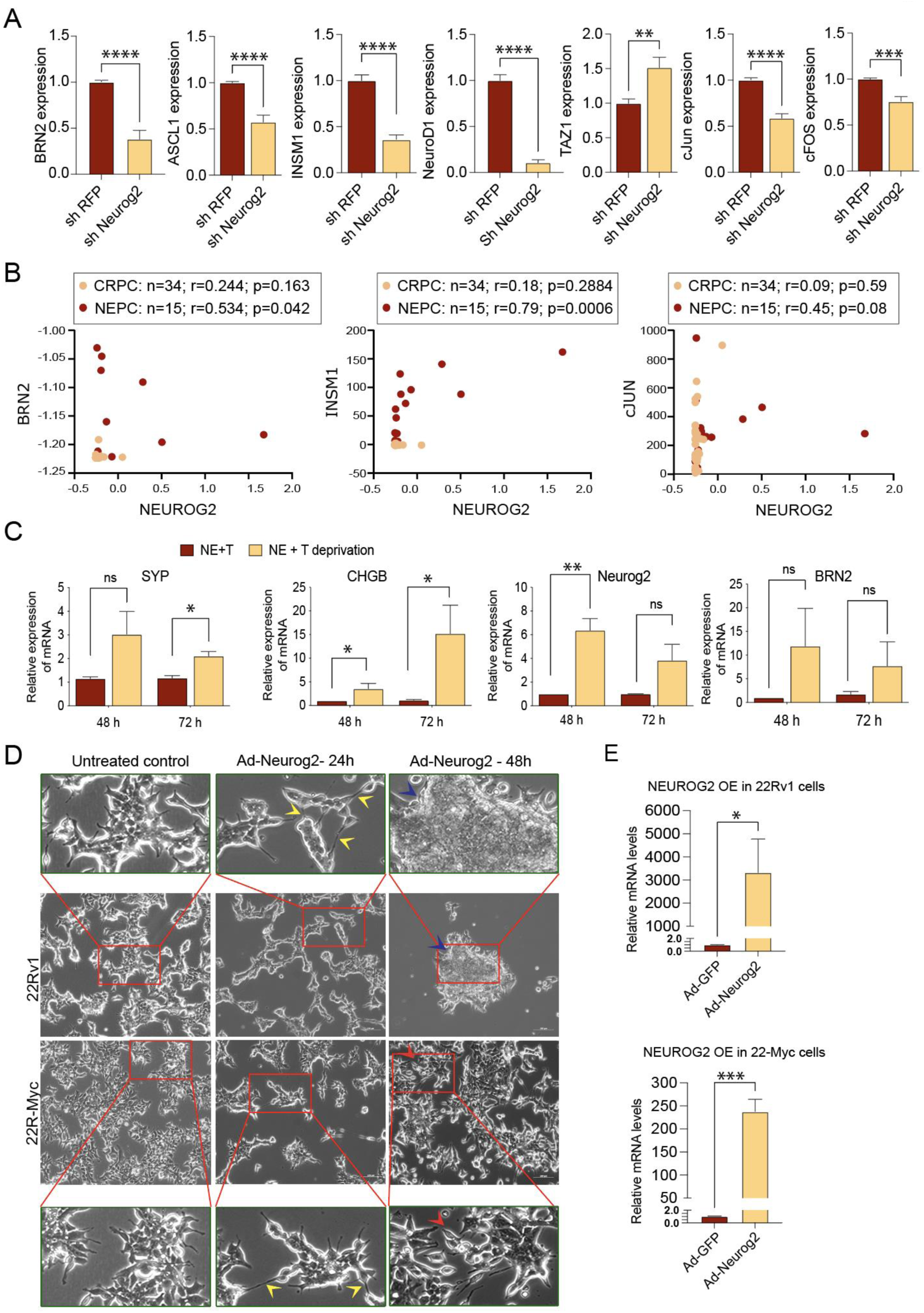
Crosstalk of Neurog2 with proneural transcription factors. (A) Knockdown of Neurog2 in LASCPC-01 cells downregulates *BRN2 (POU3F2), ASCL1, INSM1, NEUROD1, C-FOS, and C-JUN*, while upregulating *TAZ1*. Data presented as mean ± SE; n≥ 3; standard ‘t’ test; ** p<0.01; *** p<0.001, **** p<0.0001. (B) Expression correlation between Neurog2 and BRN2, INSM1 and cJUN in CRPC and NEPC samples from Beltran et al. (Nat. Methods, 2016). (C) *SYP*, *CHGB, NEUROG2 and BRN2 (POU3F2)* levels in norepinephrine (NE) supplemented but testosterone (T) withdrawn LNCaP cultures (at 48h and 72h) compared to NE+T supplemented cultures at the corresponding time points. Data presented as mean ± SE; n≥ 3; standard ‘t’ test; *p<0.05; ** p<0.01. (D) Ad-Neurog2 transduction and associated Neurog2 overexpression (OE) in 22Rv1 cells promotes cell-cell interaction through longer neurite-like extensions *(24h-yellow arrows)* and later induces clustered growth characteristics *(48h-blue arrows)*. Ad-Neurog2 transduction and associated Neurog2 OE in 22-MYC cells, while promoting cell-cell contact through neurite-like extensions *(24h-yellow arrows),* maintains the aggressive growth characteristics, including rapid filling of the growth surface *(48h-red arrows)* (scale bar, 100µm). Enlarged insets *(green bordered)* show the details of the cell phenotypes. (E) qRT-PCR shows the levels of Neurog2 mRNA in Ad-GFP and Ad-Neurog2 transduced 22Rv1 (top) and 22-MYC (bottom) cells. (Data presented as mean ± SE; n≥ 3; standard ‘t’ test; *p<0.05; *** p<0.001.

The AP-1 transcription factor complex was recently shown to promote CRPC, a subtype of PC that precedes NEPC^62^. Therefore, we next examined the expression of *c-JUN* and *c-FOS*, transcription factors that form the AP-1 transcription factor complex, in *NEUROG2* knockdown LASCPC-01 cells ^63,64^. Interestingly, we found that both *C-JUN* and *C-FOS* were downregulated after *NEUROG2* knockdown, further confirming that Neurog2 may be a potential target for tackling NEPC development **(Figure 5A).** These results were further corroborated by correlation analyses between the expression of BRN2, INSM1, c-JUN, and Neurog2 in patient samples **(Figure 5B)**. Notably, although the correlation appeared poor in CRPC (r=0.244, p=0.163 for Neurog2 vs. BRN2; r=0.18, p=0.28 for Neurog2 vs. INSM1; r=0.09, p=0.59 for Neurog2 vs. cJUN), it improved significantly in NEPC (r=0.534, p=0.042 for Neurog2 vs. BRN2; r=0.79, p=0.0006 for Neurog2 vs. INSM1; r=0.45, p=0.08 for Neurog2 vs. cJUN), indicating that Neurog2 may be a critical regulator of NEPC **(Figure 5B)**

Suppression of androgen signaling by new-generation anti-androgen therapies is a risk factor for NEPC development^4,7^. We also recently showed that neurosignaling from the sympathetic neurotransmitter norepinephrine (NE) promotes NED, a hallmark of NEPC^30^. Importantly, prostate tumors in patients receive an adequate supply of NE from sympathetic nerves ^30,65,66^. During anti-androgen therapies, androgen signaling is severely impaired, whereas intact neurosignaling is maintained. To simulate this scenario, we cultured AR^+^ LNCaP cells for two weeks in testosterone- and NE-supplemented media, and then withdrew testosterone from the cultures. As expected, testosterone depletion increased the levels of the NED markers SYP and CHGB, indicating facilitation of NEPC characteristics **(Figure 5C)**. Interestingly, this was accompanied by concomitant increases in *NEUROG2* and *BRN2* levels. These results further substantiate the potential role of Neurog2 as a master regulator of neuroendocrine plasticity.

Next, we examined the morphology of 22Rv1 and 22-MYC cells after transduction with Ad-Neurog2. Interestingly, both 22Rv1 and 22-MYC cells elongated their neurite-like extensions after Neurog2 overexpression, showing enhanced neuroendocrine characteristics **(Figure 5D and E)**. These cells also appear to form intercellular connections using neurite-like extensions. 22Rv1 cells later grew in large clusters, whereas 22-MYC cells maintained their usual growth characteristics by spreading throughout the growth surface while maintaining intercellular connections with their neurite-like extensions **(Figure 5D and E)**. Overall, the changes in the growth pattern of Neurog2 overexpressed 22Rv1 and 22-MYC cells indicated that Neurog2 induces diverse cellular fates in the presence and absence of N-Myc. Similarly, the effect of Neurog2 in inducing larger neurite-like extensions and maintaining them in N-Myc-overexpressing cells further indicated the potential of Neurog2 to promote neuroendocrine plasticity in N-Myc-overexpressing PC cells.

### Neurog2 knockdown suppresses orthotopic NEPC tumors

To further assess the relevance of Neurog2 in NEPC, we examined the effect of Neurog2 knockdown in *in vivo* orthotopic NEPC models generated using LASCPC-01 and 22-MYC cells **(Figure 6A)**. These experiments demonstrated significant suppression of NEPC tumor growth in response to *NEUROG2* silencing compared to control tumors established by matching shRFP-transduced cells **(Figure 6B)**. These findings confirm that Neurog2 has strong potential as a therapeutic target for treating patients with *MYCN* overexpressing NEPC tumors.

**Figure 6:**
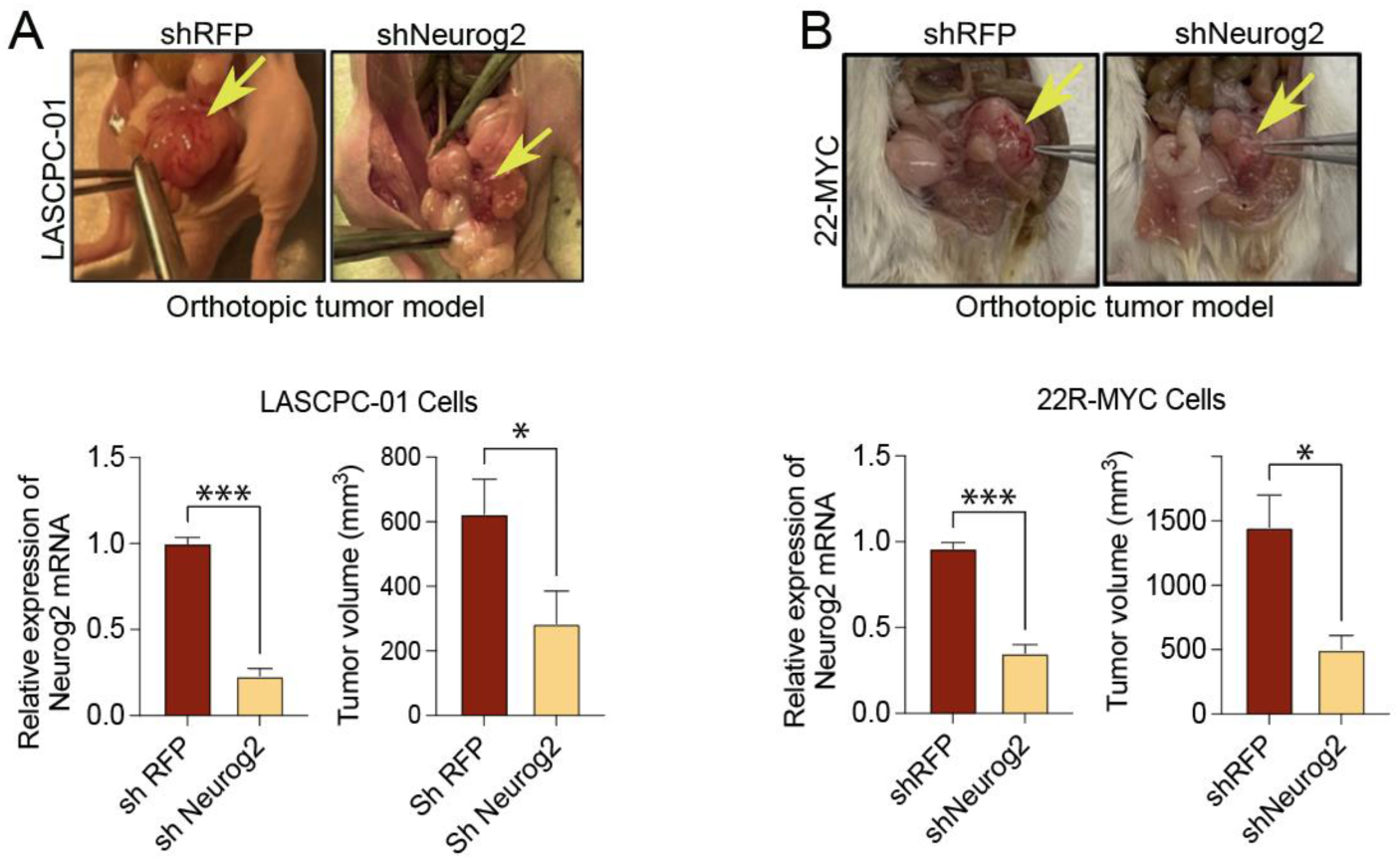
Neurog2 knockdown suppresses orthotopic tumors in NEPC models: (A) *(Top)* Representative images of orthotopic xenograft tumors induced by shRFP or shNeurog2 treated LASCPC01 cells; *(Bottom left) NEUROG2* knockdown in shNeurog2 treated LASCPC-01 cells; *(Bottom right)* quantification of tumor sizes. (B) *(Top)* Representative images of orthotopic xenograft tumors induced by shRFP or shNeurog2 treated 22-MYC cells; *(Bottom left) NEUROG2* knockdown in shNeurog2 treated 22-MYC cells; *(Bottom right)* quantification of tumor sizes. For A and B, data presented as mean ± SE; n≥ 3; standard ‘t’ test; *p<0.05; *** p<0.001.

## Discussion

Recent advances in high-quality research have paved the way for the better management of PC. Nevertheless, the most aggressive subtype of PC, NEPC, lacks efficient treatment options, which impedes disease management ^4,10,67^. Although several molecular alterations such as *RB1* loss, *TP53* loss, and *MYCN* amplification have been recognized to promote NEPC, the lack of efficient ways to directly target these molecules for therapeutic purposes hinders the development of NEPC treatment. To circumvent the complications associated with direct target modulation, we explored SDL interactions of *MYCN* by performing two independent genome-wide genetic screens using pooled shRNA and sgRNA/CRISPR/Cas9 systems. The SDL approach is a well-established method for identifying optimal therapeutic targets in a completely unbiased manner, as demonstrated by our own and previous work^17–19,21,68,69^. Using this approach, we shortlisted and preliminarily validated a few potential therapeutic targets for *MYCN* driven NEPC. While our specific focus on N-Myc^+^ NEPC may limit the applicability of our findings to selected patients, it should help in the development of effective personalized treatments for those harboring high N-Myc levels in tumors.

In this study, we conducted two independent assays to evaluate cell viability and colony formation using two different N-Myc^+^ cell lines. Additionally, we assessed the effects of the target intervention on the NED features of N-Myc^+^ cells to validate the initial set of two SDL hits identified in our screens. This strategy ensures that the optimal hit progresses towards mechanistic studies and *in vivo* assessments. Ultimately, Neurog2 PTF attracted our attention given that its inhibition selectively inhibited N-Myc^+^ cell lines and suppressed their NED signatures. Our experiments also confirmed that Neurog2 is a safe target that is not essential for normal cells, as it is not expressed in most normal adult tissues and its loss had no influence on normal cell viability. This agrees well with previously published observations showing that the functions of Neurog2 are mostly confined to developmental stages, particularly contributing to neurogenesis and maturation^47,70–73^. According to the SDL concept, inhibition of Neurog2 should preferentially eliminate N-Myc^+^ cells without affecting cells that lack N-Myc expression. Taken together, our *in vitro* validation experiments demonstrated that this is indeed true for the N-Myc-Neurog2 axis.

Neurog2 role in neurogenesis involves inducing transcriptional reprogramming in precursor cells to differentiate them into mature neurons and neuron subtypes^70^. Hence, the abundant expression of Neurog2 in N-Myc^+^ PC cells displaying NED characteristics may indicate its active role in promoting transcriptional reprogramming in PC cells for terminal transition to NEPC. Supporting this argument, the potential of Neurog2 to induce the transdifferentiation of adult cells into neuronal types has been previously demonstrated^74,75^.

Neurog2-dependent cell differentiation at developmental stages is driven by epigenetic modifications that promote chromatin accessibility at proneural genetic elements ^74,76,77^. As *NEUROG2* knockdown resulted in the downregulation of *BRN2*, *ASCL1*, and *NEUROD1* PTFs and upregulation of *TAZ1*, we expected a potential role for Neurog2 in modulating the heterochromatin landscape in N-Myc^+^ cells, favoring NEPC transition and maintenance. In addition, previous evidence suggests that epigenetic signatures consistent with NEPC favor chromatin conformations that promote access of Neurog 2 to its target genes ^78^. These findings, coupled with previous observations of Neurog-2’s capacity to transform adult cells into the neuronal phenotype, implicate Neurog-2 reactivation as a critical factor contributing to the transdifferentiation of PC into NEPC ^79^. Of the most important to note, Brn2, Ascl1 and NeuroD1 PTFs have been previously reported to promote NEPC, and hence, the collective downregulation of these PTFs downstream of Neurog2 loss observed in our investigation strongly suggests that Neurog2 may be a master regulator of PTFs, and hence, PTF-dependent NEPC. Moreover, Taz1 has been shown to suppress cellular transdifferentiation^80,81^; hence, the inverse relationship between Neurog2 and Taz1 captured in our experiments further supports a central role for Neurog2 in promoting NED characteristics in N-Myc^+^ PC cells.

With respect to profiling Neurog2 as a therapeutic target for NEPC, there is precedent for targeting transcription factors to treat NEPC. For example, in addition to the efforts of targeting N-Myc in NEPC, Nouruzi et al., showed that inhibiting the PTF Ascl1 in NEPC results in widespread epigenetic changes modulated by polycomb repressive complex-2 (PRC2) ^82^. These changes apparently reversed the neuroendocrine phenotype to a more typical luminal PC phenotype and laid the groundwork around PTFs in developing potential therapies reversing the transformation of CRPC to NEPC ^82^. Therefore, targeting Neurog2 in NEPC may offer significant therapeutic promise owing to its close association with other NEPC-related PTFs. Moreover, we discovered that Neurog2 may modulate the levels of other transcription factors such as c-Jun and c-Fos, which are linked to tissue growth. Consistent with this finding, Neurog2 knockdown effectively suppressed *MYCN*-driven NEPC tumors, highlighting the potential of Neurog2 as a therapeutic target for NEPC. Based on our experimental results, we anticipate that Neurog2 inhibitors could either revert NEPC tumors to more clinically manageable prostate cancer subtypes or serve as prophylactic agents to prevent treatment-induced NEPC. In this study, we tested Neurog2 inhibition at the preclinical level using genetic depletion approaches due to the lack of effective small-molecule inhibitors. Additional indirect approaches, such as forced induction of MTGR1, a transcriptional repressor, and feedback inhibitor of Neurog2, may be worth exploring ^72^. Another potential strategy is to inhibit specific binding of Neurog2 to its heterochromatin landscape. Further mechanistic insights into how Neurog2 orchestrates other PTFs in N-Myc^+^ NEPC could reveal additional effective intervention points.

In conclusion, understanding the intricate molecular mechanisms underlying NEPC progression and therapy resistance is crucial for the development of effective targeted therapies. In this study, leveraging the genetic concept of SDL in the context of the major genomic alteration of *MYCN* overexpression prevalent in NEPC revealed a repertoire of new potential therapeutic targets. Neurog2, one of the targets studied in more detail, shows significant promise as a focal point for developing conceptually novel and effective therapies for treating patients with N-Myc^high^ NEPC tumors.

## Acknowledgements

We sincerely thank the Rickman lab for kindly providing the 22Rv1 and 22Rv-1 N-Myc cell lines. We also thank Dr. Michael Cox for his kind contribution to BPH1, BHPrE1, NHPrE1, and PNT1B cell lines. Financial Support: This work was supported by operating grants from the Canadian Institutes of Health Research (PJT-156401; PJT-156309) to F.J.V and A.F; the Canadian Foundation for Innovation (CFI-33364) and Saskatchewan Cancer Agency operating grants with funds donated to the Cancer Foundation of Saskatchewan to F.J.V. F.S.V. is supported by the College of Medicine, U of S. H.E. was supported by the CoMGRAD award, and the U of S. J.D.W.P. is supported by grant 1160056 from the Cancer Research Society. V.M. is supported by the Canada Graduate Scholarship - Doctoral from Canadian Institutes of Health Research (FBD- 187665) as well as the Health Sciences Graduate Scholarship from the College of Medicine at the University of Saskatchewan. This work was also supported by the CoMBRIDGE award (A.K, F.J.V, A.F), funding from the Saskatchewan Health Research Foundation (Establishment Grant), and Prostate Cancer Fight Foundation and Ride for Dad to AK.

## Author contributions

Conceptualization: A.K., A.F., F.J.V.

Investigation: P.W., J.D.W.P., F.S.V., N.J., B.C., J.P., Y.Z., H.D., L.N., A.G., K.H.G., H.E., and K.K.B.

Writing – Review & Editing: All authors.

Funding Acquisition: A.K., A.F. and F.J.V.

## STAR Methods

### Resources availability

#### Lead contact

Further information and requests for reagents and resources should be directed to and will be fulfilled by the lead contact, Franco Vizeacoumar.

#### Materials availability

This study did not generate new unique reagents.

#### Data and code availability

The data relevant to this study were deposited in the Gene Expression Omnibus (GEO) database. The microarray shRNA screening data can be found under accession number <u>GSE255592</u>, and the CRISPR screening sequencing data are available under accession number <u>GSE255591</u>. The comprehensive dataset was accessible using super series <u>GSE255593</u>. All other data reported in this paper will be shared by the lead contact upon request. This paper does not report original code. Any additional information required to reanalyze the data reported in this paper is available from the lead contact upon request.

### Experimental model and study participant details

#### Cell lines and cell culture

LASCPC-01 cells (ATCC® CRL-3356) and LNCaP cells (ATCC® CRL-1740) were obtained from ATCC (Manassas, VA, USA) and cultured in modified HITES medium as previously described^49^. The isogenic cell line pairs 22Rv1 and 22Rv1-N-Myc (22-MYC) were provided by the Rickman Lab (Cornell Medical College). The non-malignant cell lines BPH1, BHPrE1, NHPrE1, and PNT1B were kindly provided by Dr. Michael Cox at the Vancouver Prostate Center. 22Rv1, BPH1, PNT1B, and LNCaP cells were maintained in RPMI-1640 containing 10% FBS and 1% penicillin/ streptomycin. HEK293T cells were cultured in DMEM containing 10% FBS and 1% penicillin/ streptomycin. NHPrE1 and BHPrE1 were cultured in DMEM/F12 media with 5% FBS, 1% insulin-transferrin-selenium-X, 0.4% Bovine Pituitary Extract, 10 ng/L EGF, and 1% penicillin/streptomycin. Cell lines were incubated at 37°C in 5% CO2.

#### Orthotopic prostate cancer model

Mice were housed 5 per cage at 23-25°C in a humidity-controlled colony room, maintained on a 12h light/dark cycle (08:00 to 20:00 light on), with standard food and water provided ad libitum and environmental enrichments. All animals were handled in accordance with the approved protocols by the University of Saskatchewan Animal Research Ethics Board (AREB). Mice used in this study were nude (Crl:NU(NCr)-*Foxn1^nu^*) for LASCPC-01 experiments and NSG (NOD.Cg-Prkdscidll2rg) for 22-MYC experiments, and littermates within the same cage were randomly allocated to experimental or control groups.

All animal experiments were approved by the Animal Ethics Committee of the University of Saskatchewan. Adult male nude mice weighing 20-25 g were used in this study. Five mice were allocated to each experimental group. Both 22-MYC and LASCPC-01 cells were used for tumor induction. To perform orthotopic injection of cells into the prostate, the mice were anesthetized using isoflurane and buprenorphine (0.05 mg/kg). A small incision was made in the lower abdomen, and the urinary bladder was exposed. Orthotopic injection was performed by injecting 200,000 cells in 1:1 media and Cultrex basement membrane extract (R&D Systems) into the exposed prostate. The incision was sutured, and the animals were allowed to survive for four weeks. Tumor sizes were recorded using a Vernier caliper *(for LASCPC-01 experiments)* or digital calipers *(for 22-MYC experiments)* at experimental termination. Tumor sizes were calculated using the standard formula LxW^2^/2, where L is the length and W is the width of the tumor.

### Methods details

#### Lentivirus production

HEK-293T cells were used to produce lentiviral particles expressing short hairpin RNA (shRNA) or sgRNA. This process involved packaging 5400 ng of psPAX2 and 600 ng of pMD2.G plasmids in conjunction with 6000 ng of target gene plasmids. Next, we used 10 cm plates to transfect the cells with a medium consisting of 540 µL Opti MEM (Gibco Life Technologies) and 36 µL of X-treme GENE DNA Transfection Reagent (Roche, Mississauga, ON, Canada). After 18 h, the medium was replaced with DMEM containing 2% bovine serum albumin (MilliporeSigma). Supernatants containing viral particles were collected 24 and 48 h post-transfection after centrifugation at 185 × G for 5 min. The supernatants were stored at -80 C for subsequent use. Stable 22-Rv1-Cas9 and 22-MYC-Cas9 cells were generated by transducing the cells with Cas9-blasticidin lentivirus and 8 µg/mL polybrene (MilliporeSigma). The medium containing blasticidin (4µg/ml) was changed every 2-3 days (for 10-12 days) until the non-transduced cells were eliminated.

#### Pooled shRNA and CRISPR/Cas9 screening and data deconvolution

Lentiviral transduction of 22Rv1 and 22-MYC cells with shRNA or sgRNA libraries at 0.3 MOI was done as previously described^18–20^. Briefly, 24 h post-transduction, the cells were treated with 2 ug/ml of puromycin for 48 h. Puromycin-selected cells were then passaged for 12 d, and the cells were collected at three time points (Day 0, 7, and 13 for shRNA and Day 0, 6, and 12 for CRISPR screens). For shRNA screening, microarray analysis was performed, and deep sequencing was performed for the described previously^18–20^. Genomic DNA was extracted at each time point and the shRNA sequences were amplified. The amplified hairpins were digested with Xhol and purified for probe hybridization. FASTQs were aligned to library sequences using MAGeCK software (version 0.59) for CRISPR/Cas9 screens^83^. Prior to alignment, the FASTQC package (version 0.12.1) was used for quality assurance. These read counts were then cyclic-loss-normalized before computing the fitness score. shRNA- or sgRNA-weighted differential cumulative change (WDC_h/g_) between 22RV1 and 22-MYC was calculated at each corresponding time point using the following equation:

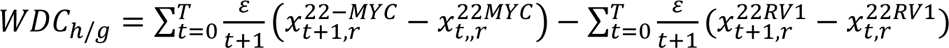

where 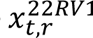 is the normalized signal intensity of 22RV1 cells at time point *t* ∈ (0, . . *T*) in replicates *r* ∈ (1. . *N*). Similarly, 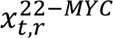 was detected in the 22-MYC cells. *ε* is a constant that improves the weighting of shRNA or sgRNA hits the earlier the gene of interest drops. The WDC gene level WDC_gene_ was calculated as the average of the top two shRNA/sgRNA with the lowest value for that gene using the following equation:

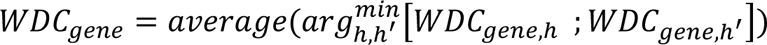

Significant differences between shRNA or sgRNA and their corresponding genes in 22RV1 and 22-MYC cells were determined by Student’s t-test in combination with the permutation test p-value by estimating the frequency of randomized, shuffled WDC with lower values in comparison to the observed gene level WDC value^20^. Bayesian analysis of gene essentiality was used to evaluate screen performance^84^.

#### Pathway enrichment analysis

Genes that showed significant WDC_gene_ values from both CRISPR and shRNA screens were combined for comprehensive pathway enrichment analysis. This analysis was conducted using Gene Set Enrichment Analysis (GSEA) software version 4.2.2^85^. The focus was on identifying enriched gene ontology biological processes (version 2023.2). Pathways were considered enriched if they had a false discovery rate (FDR) of less than 0.001 and a pathway similarity cutoff greater than 0.5. These enriched pathways were imported into the enrichment map visualization plug-in for further examination^86^. The results were subsequently exported to Cytoscape for detailed network visualization^87^. To provide a clearer understanding of the contributions of each screen, the pathways were segmented and visualized as pie charts. These charts illustrate the proportion of genes identified from the CRISPR screen versus those identified from the shRNA screen, offering a visual representation of the data distribution and relative impact of each screening method on the identified pathways.

#### Differential expression analyses of CRPC and NEPC patient samples

Published data from Beltran et al.,^35^, which included 34 castration-resistant PC samples with adenocarcinoma features (CRPC-Adeno) and 15 castration-resistant PC samples with neuroendocrine features (CRPC-NE), were used to identify genes relevant to neuroendocrine features that overlapped in our shRNA and CRISPR screens. From the initial pool of 1120 genes identified by both screening methods, 323 genes were found to be significant (p<0.05) based on the non-parametric Wilcoxon rank-sum test. The expression levels of these 323 genes were normalized across all CRPC-adeno and CRPC-NE samples. Subsequent clustering was performed using Pearson’s correlation to analyze the expression patterns. This clustering approach allowed for the comparison of gene expression profiles between CRPC-Adeno and CRPC-NE samples, highlighting distinct expression signatures associated with neuroendocrine differentiation.

#### Expression analysis of TCGA datasets

To investigate the differential expression of neuronal plasticity genes in lung cancer subtypes, we utilized data from The Cancer Genome Atlas (TCGA), which includes samples of lung adenocarcinoma (LUAD) and lung squamous cell carcinoma (LUSC). Tumor patients were categorized into two groups of high and low expression based on the median expression levels of four neuroendocrine genes in normal tissues: *MYCN, synaptophysin (SYP), chromogranin A (CHGA)*, and neuron-specific enolase (*NSE)*. We focused on 17 genes related to neuronal plasticity. The expression levels of these genes were compared between the two groups with high and low neuroendocrine gene expression. The statistical significance of the differences in neuroendocrine gene expression between the two groups was assessed using a one-tailed unpaired t-test performed in GraphPad Prism. This test allowed us to determine whether there were significant differences in the expression levels of neuronal plasticity genes between the high- and low-expression groups.

#### Neurog 2 overexpression studies

Adenoviral particles carrying the human Neurog 2 plasmid were purchased from Vector Biolabs (SKU-ADV-216616). 5×10^5^ 22Rv1 and 22-MYC cells were seeded in 6-well plates containing 1mL culture media. After 24 h, the medium was replaced with 1mL fresh media, and the cells were infected with adenoviral particles (50 MOI) in the presence of 8 µg/mL polybrene (MilliporeSigma; TR-1003). Twenty-four hours after transduction, the media was removed, the wells were washed with PBS, and fresh media was added. Brightfield images were taken using a Zeiss Axio Observer under 10x magnification at 24 and 48 h after infection.

#### qRT-PCR

RNA was isolated and purified using a PureLink RNA isolation kit (ThermoFisher) according to the manufacturer’s protocol. RNA was reverse transcribed to cDNA using a cDNA synthesis kit (Applied Biosystems) in a master cycler. qRT-PCR was carried out using the StepOne Real-Time PCR System (Applied Biosystems) and PowerUp^TM^ SYBR^TM^ Green Master Mix (Applied Biosystems). The primers used to amplify cDNAs are listed in Supplementary Table S1. All reactions were performed in triplicate. All target genes were internally normalized to the housekeeping genes (GAPDH, RPLP, and actin).

#### Colony formation and cell viability assays

For colony formation assay, the control and target shRNA- or sgRNA treated 22Rv1 and 22-MYC cells were seeded (1000 cells/well) in a 6-well plate and incubated at 37 °C and 5% CO_2_ for 12 d, replacing the media every 3 days. The plates were then washed with PBS three times, fixed with 4% formalin solution, and stained with 0.5% crystal violet. Images of the colonies were captured using a Bio-Rad Imager, and the colonies were quantified using ImageJ software. For the cell viability assay, metabolically active cells were quantified using either a resazurin reduction assay or a CCK8 assay after the knockdown of target genes. Briefly, LASCPC-01 cells or non-malignant BPH1, BHPrE1, NHPrE1, and PNT1B cells were transduced with shNeurog2, shIsl1, or shRFP, followed by puromycin selection for 48 h. Puromycin-selected cells (1000 cells) were seeded into 96 well plates. A 1:10 dilution of resazurin or 10 µL of CCK8 reagent was then added to each well of a 96-well plate and incubated for 4 h at 37 °C and 5% CO_2_. Fluorescence intensity was measured at 590 nm (resazurin) or 450 nm (CCK8) using a Varioskan LUX microplate reader.

#### Correlation analysis of gene expression

Published data from Beltran et al. ^35^ was utilized to perform a comparative analysis of gene expression. The expression levels of SYP, CHGA, and NSE were compared to those of Neurog2. Similarly, the expression levels of BRN2, INSM1, and cJUN were also compared with those of Neurog2. Spearman rank correlation was used to calculate the correlation coefficient (r) and determine the statistical significance of these relationships.

### Quantification and statistical analysis

GraphPad prism software was used for all tests. Unless otherwise specified, analyses were done using standard ‘t’ tests, one-way ANOVA (with Tukey’s multiple comparisons test) or two-way ANOVA (with Sidak’s multiple comparisons test). The details of the statistical analysis performed, including the specific test employed, the number of samples considered, and the level of significance achieved, are presented in more detail in the corresponding figure legends.

## Supplemental information

**Supplementary Figure S1:**
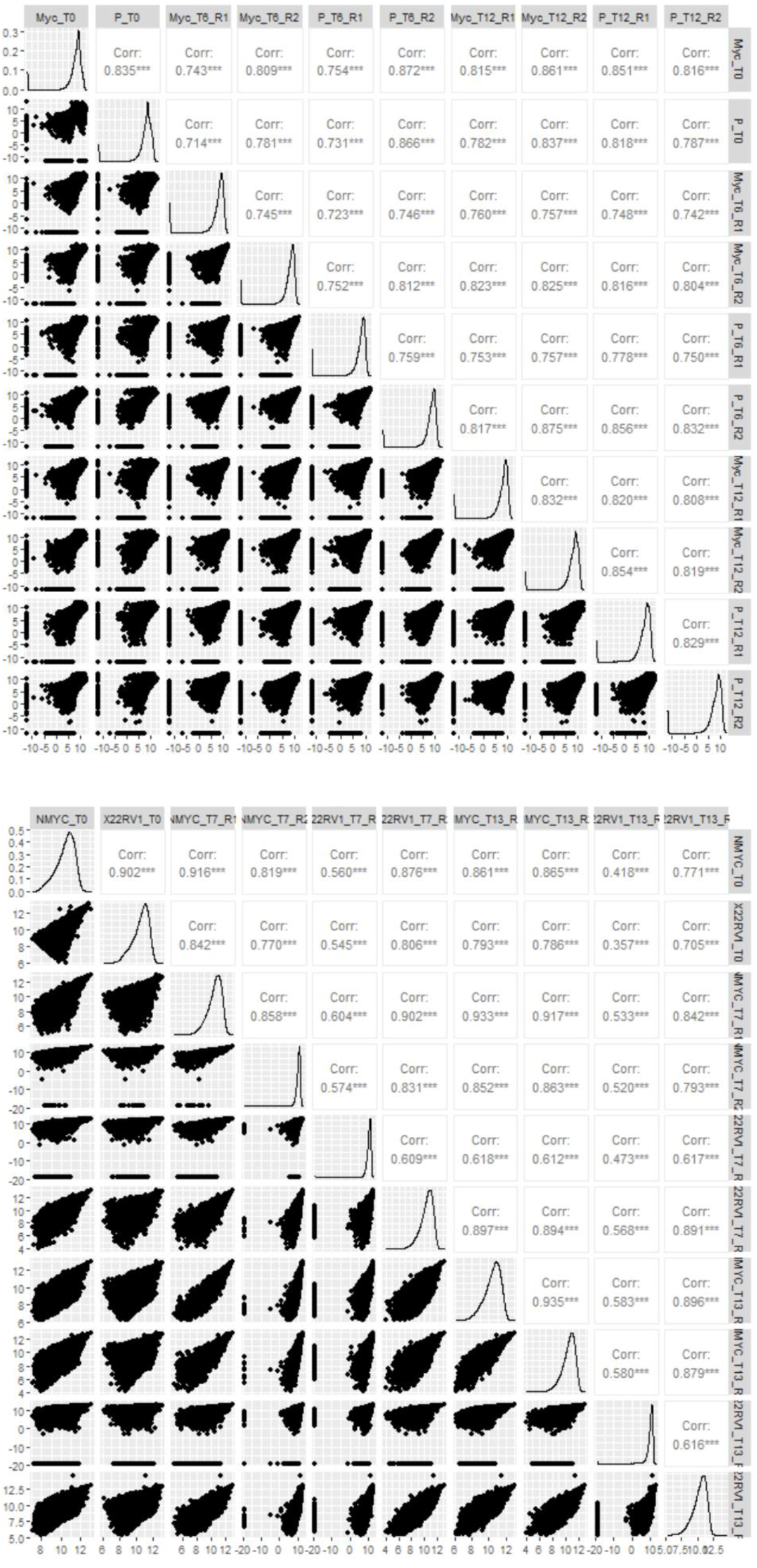
Correlation of replicates in shRNA (top) and CRISPR screens (bottom).

**Supplementary Figure S2:**
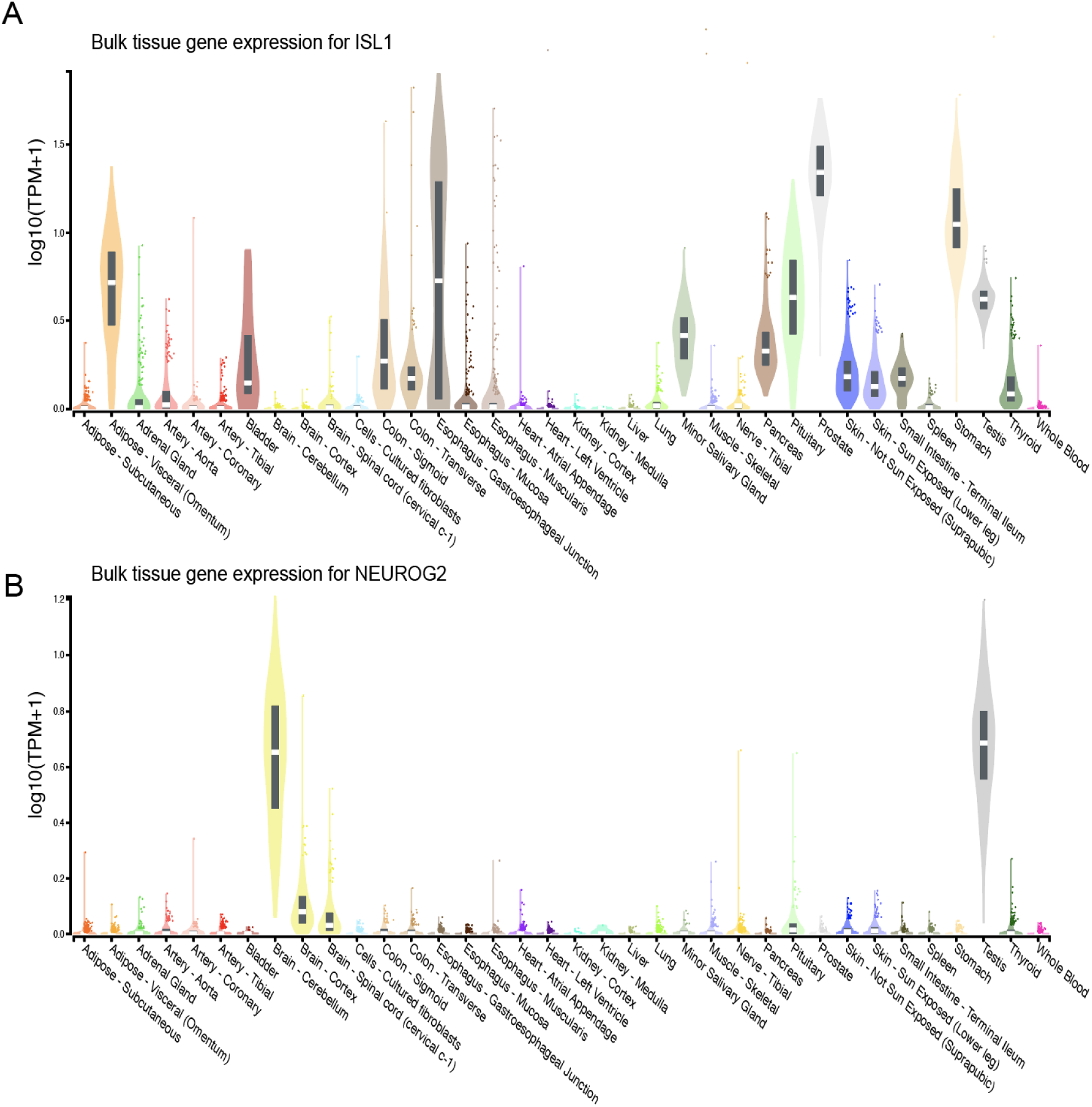
Gene expression data of *ISL1* (A) and *NEUROG2* (B) in normal human tissues derived from GTEx portal.

**Supplementary Figure S3:**
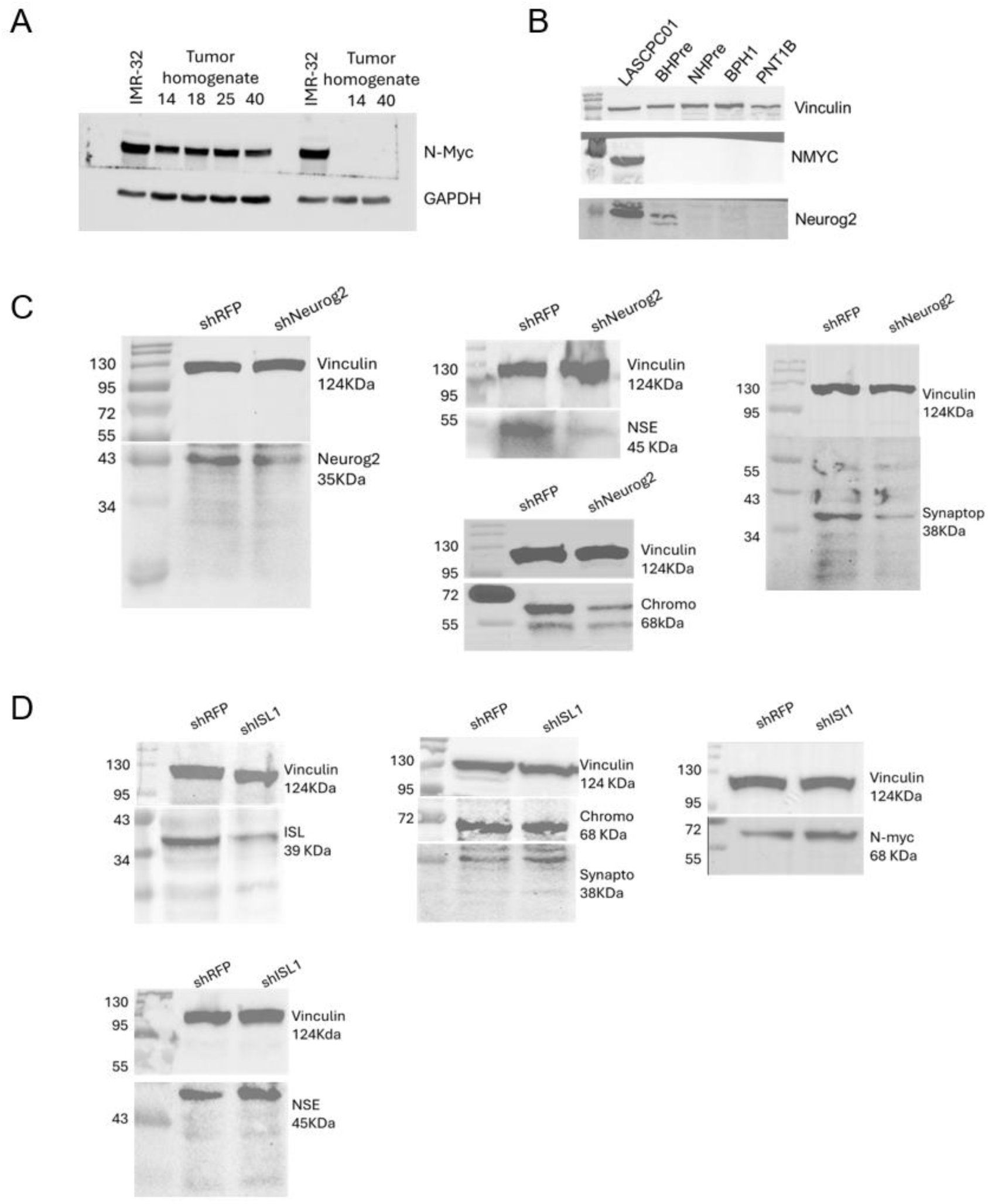
Original Western blots relating to Figure 1 (A), Figure 3 (B) and Figure 4 (C and D).

**Supplementary Table S1:** qPCR Primers

**Supplementary Table S2:** List of 553 genes that are SDL to NMYC identified from CRISPR/Cas9 screen

